# Telomerase mRNA-Lipid nanoparticles attenuate neuroinflammation after traumatic brain injury in mice

**DOI:** 10.64898/2025.12.18.694748

**Authors:** Goknur Kara, Morgan Holcomb, Anjana Tiwari, Hannah Flinn, Trinity Eimer, Austin Marshall, Marissa Burke, Peter Park, Karem Court, John P Cooke, Biana Godin, Sonia Villapol

## Abstract

Traumatic brain injury (TBI) is a leading cause of chronic neurological disability, yet no disease-modifying therapy exists. Emerging evidence indicates that TBI activates cellular aging programs, including telomere erosion and persistent inflammation, that contribute to progressive neurodegeneration. Telomerase reverse transcriptase (TERT) preserves telomere homeostasis and provides cytoprotective effects in the central nervous system, but has not been therapeutically targeted after TBI. Here, we developed an mRNA nanotherapy consisting of mouse TERT mRNA encapsulated in lipid nanoparticles (mTERT-LNPs) and evaluated it in a controlled cortical impact model of moderate TBI. We first established that TBI transiently disrupts TERT biology, with reduced cortical TERT mRNA and shortened telomeres at 3 days post-injury (dpi), followed by partial recovery by 14 dpi. mTERT-LNPs were well tolerated in vitro and in vivo. Following intravenous delivery in the acute post-injury window, LNPs localized to the injured brain and displayed expected peripheral biodistribution. A single systemic dose increased cortical TERT mRNA and protein and partially restored telomere length at 3 dpi. TERT mRNA delivery significantly reduced Iba1+ microglial activation and suppressed pro-inflammatory cytokines, with modest increases in anti-inflammatory markers. Systemically, mTERT-LNPs lowered serum C-reactive protein and malondialdehyde, indicating reduced peripheral inflammation and oxidative stress, without adverse effects on body weight or peripheral organ histology. Several outcomes showed sex-dependent patterns. Collectively, these data provide the first in vivo evidence that telomerase therapy can modulate telomere biology and neuroinflammation after TBI, supporting mRNA-LNP-mediated TERT restoration as a scalable, mechanistically grounded strategy for disease modification in TBI and related disorders.

**Significant statement:** Traumatic brain injury (TBI) can initiate progressive brain changes that worsen long after the initial impact, and no therapy directly slows or prevents this decline. A key contributor may be “accelerated aging” in injured tissue, including telomere damage (protective chromosome ends) and persistent neuroinflammation driven by activated immune cells. This study is significant because it tests a targeted way to disrupt these processes by restoring telomerase reverse transcriptase (TERT), which helps maintain telomeres and cellular resilience. Using a clinically validated mRNA-lipid nanoparticle platform, a single intravenous dose increased brain TERT, partially restored telomere length, and reduced neuroinflammation and systemic oxidative stress without obvious toxicity. These findings connect telomere dysfunction to a scalable disease-modifying strategy for TBI and related neurodegenerative conditions.

## INTRODUCTION

Traumatic brain injury (TBI) is a major cause of death and long-term disability, affecting ∼69 million people worldwide annually and ∼2.8 million only in the United States (Li et al., 2021; Zhang et al., 2021). Beyond the acute insult, TBI is now understood as a chronic disorder that accelerates late-life neurodegeneration (Maas et al., 2022) and increases the risk for Alzheimer’s disease (Gardner and Zafonte, 2016), Parkinson’s disease, and chronic traumatic encephalopathy (Bielanin et al., 2024). Secondary injury cascades, including blood-brain barrier disruption, axonal degeneration, mitochondrial dysfunction, oxidative stress, and sustained neuroinflammation, drive progressive tissue loss and functional decline (Thapa et al., 2021; Navabi et al., 2024). Despite intensive clinical and preclinical efforts, no disease-modifying therapy has yet proven effective in improving long-term outcomes of TBI.

Accumulating evidence indicates that TBI also activates cellular aging programs. Telomeres, the repetitive DNA sequences that cap chromosome ends, shorten with cell division, oxidative stress, and inflammation (Aubert and Lansdorp, 2008; Maynard et al., 2015; Liu et al., 2019). Clinical and experimental studies show that telomere length (TL) is reduced after TBI and may serve as a biomarker of injury severity and outcome (Smith et al., 2013; Eitan et al., 2014; Wright et al., 2018; Ng and Lee, 2019; Zheng et al., 2022; Martha et al., 2023). For example, repetitive mild TBI in rodent models induces significant telomere shortening within days of injury, coinciding with heightened oxidative stress and neuroinflammation (Wright et al., 2018). These findings suggest that telomere erosion is an active component of TBI pathophysiology.

Telomerase reverse transcriptase (TERT), the catalytic subunit of telomerase, maintains telomere homeostasis, but also exerts non-canonical, telomere-independent functions in the central nervous system. Beyond its role in telomere extension (Saretzki and Wan; Spanakis and Tsatsakis, 2025), TERT protects neurons from oxidative damage, modulates mitochondrial function, reduces toxic protein accumulation, and dampens senescence-associated inflammation while promoting neurogenesis and cognition in multiple preclinical models (Li et al., 2013; Spilsbury et al., 2015; Miwa et al., 2016; Wang et al., 2020; Yu et al., 2023; Shim et al., 2024). Telomerase activation has, therefore, emerged as a promising therapeutic strategy in neurological disease; however, telomerase-targeted interventions have not been evaluated in TBI. Recently, we demonstrated that delivery of human TERT mRNA encapsulated in lipid nanoparticles (LNPs) enhances human skin cell suspension engraftment and proliferation in a humanized mouse wound-healing model, establishing the feasibility and safety of TERT mRNA-LNP therapy *in vivo* (Chang et al., 2024). We also reported that TERT mRNA-LNPs significantly reduced radiation-induced DNA damage in human primary skin cells and tissues, enhanced DNA repair, decreased mitochondrial ROS, and lowered apoptosis, without extending telomere length during the experimental period, indicating a non-canonical role of TERT in accelerating cellular recovery from radiation (Li et al., 2025). mRNA therapeutics provide a flexible platform for the transient expression of therapeutic proteins, including targets that are otherwise considered “undruggable” (Chanda et al., 2021; Metkar et al., 2024). Advances in *in vitro*-transcribed (IVT) mRNA chemistry and LNP formulations, highlighted by the success of mRNA-LNP COVID-19 vaccines, have transformed mRNA into a clinically validated modality. Since naked mRNA is rapidly degraded in biological fluids, efficient delivery systems such as LNPs are essential to protect mRNA, promote cellular uptake, and enable controlled in vivo protein production (Chanda et al., 2021; Chang et al., 2024; Metkar et al., 2024). Among non-viral vectors, LNPs have shown the most advanced safety and efficacy profile in humans, making them attractive candidates for CNS-targeted gene therapies.

Building on prior work linking TBI to telomere attrition and on our previous demonstration of TERT mRNA-LNPs’ efficacy in a regenerative setting, we hypothesized that transient restoration of TERT expression after TBI could counteract injury-induced telomere shortening and attenuate downstream neuroinflammation. Here, we develop and characterize a mouse TERT (mTERT) mRNA-LNP formulation and test its therapeutic potential in a controlled cortical impact model of moderate TBI. We show that a single intravenous dose of mTERT-LNPs administered shortly after injury enhances cortical TERT expression, partially restores telomere length, and dampens microglial activation and pro-inflammatory cytokine production, with sex-dependent effects, while maintaining a favorable systemic safety profile. These findings position TERT mRNA-LNP therapy as a mechanistically grounded, clinically translatable strategy to mitigate secondary injury and potentially modify long-term outcomes after TBI.

## MATERIAL AND METHODS

### Synthesis of messenger RNA and lipid nanoparticles (LNPs)

Mouse TERT (mTERT) mRNA was synthesized by the Houston Methodist Research Institute (HMRI) RNA Core through IVT, with pseudouridine added to the nucleotide mix to enhance its translation and to reduce the inflammatory response caused by the mRNA, as previously described (Ramunas et al., 2015a). Firefly luciferase (Luc) mRNA was purchased from TriLink Biotechnologies (cat. #L-7602). mRNA-LNP formulations were generated using a NanoAssemblr Benchtop (Precision Nanosystems). Briefly, LNPs were formulated using a molar ratio of 8:1.5:38.5:52 for Distearoylphosphatidylcholine (DSPC, Avanti Polar Lipids, cat. #850365), 1,2-Dimyristoyl-rac-glycerol-3-methoxy polyethylene glycol-2000 (DMG-PEG 2000, Avanti Polar Lipids, cat. #880151P), cholesterol (Millipore Sigma, cat. #3667), and Dlin-MC3-DMA (MedChemExpress, cat. #HY-112251), respectively. For biodistribution studies, LNPs were fluorescently labeled with 1,2-dioleoyl-sn-glycero-3-phosphoethanolamine-N-Cyanine 5.5 (Avanti Research, cat. #810336). The lipids were dissolved in ethanol, and the mRNA in citrate buffer (100 mM, pH 5.0). The aqueous and ethanolic phases were further mixed at a 3:1 ratio at a flow rate of 10 ml min^−1^. Subsequently, dialysis was performed in PBS at 4°C for at least 12 h to eliminate residual ethanol and unbound mRNA (Chang et al., 2024). The size, polydispersity index (PDI), and zeta potential of the LNPs were measured using dynamic light scattering (DLS) with a Zetasizer NanoZS (Malvern Instruments). The mRNA encapsulation efficiency (EE) in the LNPs was measured using RiboGreen Assay (Fisher Scientific, cat. #R11490) based on the manufacturer’s protocol. Both encapsulated and unencapsulated mRNA were measured. The total mRNA (encapsulated + unencapsulated) was measured following LNPs digestion with 1% Triton X-100 to release the RNA’s contents. EE was calculated using the formula:

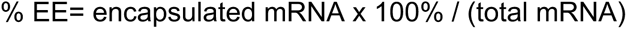

### *In Vitro* Viability in Neuronal Cell Culture

N2a (Neuro-2a, cat. # CCL-131) murine cell line was obtained from ATCC (US). The cells were cultured with Dulbecco’s modified Eagle’s medium (DMEM, Thermo Fisher Scientific) supplemented with 10% fetal bovine serum (FBS, Thermo Fisher Scientific) and 1% penicillin-streptomycin. *In vitro* biocompatibility of Luc mRNA-LNPs and mTERT mRNA-LNPs was determined using the WST-1 assay. Cells were seeded at a density of 3000 cells per well in 96-well plates and incubated overnight. Subsequently, they were treated with mRNA-LNPs at 0, 0.5, 1, and 2 µg/mL of mRNA, followed by incubation for 24 and 48 h (n = 5). WST-1 assay reagent (Millipore Sigma, cat. #12352200) was added at the end of the incubation period, and the plates were incubated for 2.5 h. Absorbance was measured at 490 nm, and the values were normalized to the untreated control.

### Mice and TBI model

6-month-old male and female C57BL/6J mice from Jackson Laboratories (Bar Harbor, ME, US) were housed at the HMRI animal facility. The mice were kept under a 12 h light-dark cycle and provided *ad libitum* access to food and water. All animal experiments adhered to approved protocol as specified by the Institutional Animal Care and Use Committee (IACUC) at HMRI. These procedures were conducted in accordance with established institutional guidelines and regulations. All mice were anesthetized with isoflurane during surgeries, starting at 3% for induction and maintained at 1.5-2%. We used an electromagnetic Impact One stereotaxic impactor (Leica Biosystems, Buffalo Grove, IL, USA) to induce moderate TBI in the left hemisphere of mice, targeting the primary motor and somatosensory cortices (Villapol et al., 2017; Holcomb et al., 2025; Soriano et al., 2025). The impact site was located 2 mm lateral and 2 mm posterior to Bregma using a flat impact tip with a diameter of 3 mm. The impact was delivered at a velocity of 3.2 m/s and a depth of 1.5 mm.

### Biodistribution of mRNA-LNPs using IVIS

Brain accumulation and biodistribution of mRNA-LNPs were assessed in male and female mice with TBI using an *in vivo* imaging system (IVIS). For these studies, Cy5.5-labeled LNPs containing Luc-mRNA were used to track the LNP distribution and protein expression. Mice received an intravenous injection of either Luc-Cy5.5-LNPs or mTERT-Cy5.5-LNPs 30 min post-injury, with a dose of 1-3 mg/kg of mRNA. 24 h after TBI, mice treated with Luc-Cy5.5-LNPs were intraperitoneally injected with 150 mg/kg luciferin immediately before IVIS analysis (Perkin Elmer IVIS). Mice were then euthanized, and their organs, including the brain, heart, lungs, spleen, liver, and kidneys, and blood were collected for imaging. Fluorescence (ex = 640 nm, em = 720 nm) and bioluminescence were measured. Data analysis was performed using the Living Image software.

### Telomere length (TL) analysis

We assessed TL using a relative mouse telomere length quantitative real-time PCR (qRT-PCR) assay kit (ScienCell Research Laboratories, cat. #M8908) according to the manufacturer’s instructions. Before the assay, we isolated genomic DNA from the injured brain sites of each mouse using the TRIzol Reagent (Invitrogen, cat. #15596026) protocol. We performed two qPCR reactions for each genomic DNA sample using telomere and single-copy reference (SCR) primers, along with TaqGreen qPCR master mix. The SCR primer set was used as a reference for data normalization. The PCR cycling protocol was set up as follows: an initial denaturation step at 95°C for 10 min, followed by 32 cycles of denaturation at 95°C for 20 s, annealing at 52°C for 20 s, and extension at 72°C for 45 s. The relative TL was determined by calculating the T/S ratio, which indicates the amplification of the telomere product (T) relative to the single-copy reference gene (S). The relative T/S ratio was computed using the 2^−ΔΔCt^ method, which compares the variation of each DNA sample’s T/S ratio to a reference sample (Cawthon, 2002). Measurements were performed in triplicate for all samples.

### RNA extraction and qPCR

Total RNA was extracted from the injured brain tissue using TRIzol™ Reagent (Invitrogen, cat. #15596026) following the manufacturer’s recommended protocol. One µg of isolated RNA from each sample was used as a template and reverse-transcribed into complementary DNA (cDNA) using the iScript cDNA Synthesis Kit (Bio-Rad, cat. #1708891) with the following thermal cycle conditions: 25°C for 5 min, 46°C for 20 min, and 95°C for 1 min. Relative gene expression was normalized to the housekeeping control gene, β-actin. cDNAs were amplified using SsoAdvanced Universal SYBR Green Supermix with the CFX384 Touch Real-Time PCR Detection System (BioRad), and the relative differences in gene expression were determined using the comparative threshold cycle (2^−ΔΔCt^) method.

Sequences of the primers used are as follows: Tert: F:5’-TCTCTATGAATGAGAGCAGC-3’ and R:5’-TATAGCACCTGTCACCAATC-3’; TNF-α: F:5’-CTATGTCTCAGCCTCTTCTC-3’ and R:5’-CATTTGGGAACTTCTCATCC-3’; IL-1β: F:5’-GGATGATGATGATAACCTGC-3’ and R:5’-CATGGAGAATATCACTTGTTGG-3’; IL-6: F:5’-AAGAAATGATGGATGCTACC-3’ and R:5’-GAGTTTCTGTATCTCTCTGAAG-3’; IL-18: F:5’-AAATGGAGACCTGGAATCAG-3’ and R:5’-CCTCTTACTTCACTGTCTTTG-3’; TGF-β: F:5’-GGATACCAACTATTGCTTCAG-3’ and R:5’-TGTCCAGGCTCCAAATATAG-3’; IL-10: F:5’-CAGGACTTTAAGGGTTACTTG-3’ and R:5’-ATTTTCACAGGGGAGAAATC-3’; and β-Actin: F:5’-GATGTATGAAGGCTTTGGTC-3’ and R:5’-TGTGCACTTTTATTGGTCTC-3’.

### Western Blot

Serum was obtained by centrifuging the blood samples collected from each mouse at 4000 rpm for 20 min at 4°C. Next, serum samples (1:5 v/v) were combined with 4x Laemmli sample buffer (Bio-Rad, cat. #1610747) and incubated at 100°C for 5 min. The samples were subjected to SDS-PAGE with a 4% to 15% gradient for protein separation and electro-transferred to polyvinylidene difluoride (PVDF) membranes (Bio-Rad, cat. #1620177). The membranes were blocked in 5% w/v skim milk powder in PBS-Tween 20 (PBS-T) buffer for 1 h at room temperature. The expression level of C-reactive protein (CRP) was detected using a specific CRP antibody (Proteintech, cat. #66250-1-Ig) and the corresponding HRP-conjugated secondary antibody. Immunoblots were visualized using Clarity Western ECL (Bio-Rad, cat. #1705061) in a ChemiDoc MP imaging system (Bio-Rad) and quantified with a densitometer using ImageJ.

### MDA analysis

Malondialdehyde (MDA), the end product of lipid peroxidation, was determined in serum using the MDA assay kit (Abcam, cat. #ab118970) according to the manufacturer’s instructions. The color reaction was measured at 540 nm. The MDA levels were expressed as mM.

### Brain tissue preparation and fluorescent in situ hybridization with immunohistochemical labeling

Brain samples were fixed in 4% paraformaldehyde overnight, then transferred to 30% sucrose for additional processing. Using a cryostat (Epredia Cryostar NX50, Fisher Scientific), the brains were sectioned into 16 μm slices. Sections from the frontal cortex through the dorsal hippocampus were collected at coronal planes. These sections were either mounted directly onto glass slides or kept in a free-floating cryoprotective solution containing 30% sucrose, 1% polyvinylpyrrolidone, 30% ethylene glycol, and 0.01 M PBS.

Coronal brain sections were placed on gelatin-coated glass slides (Superfrost Plus, Fisher Scientific, cat. #12-550-15) and kept at −80°C until they were used. Fluorescent in situ hybridization (FISH) was performed according to the manufacturer’s instructions using the RNAscope™ 2.5 HD Reagent Kit-RED (Advanced Cell Diagnostics, cat. #322350), as previously described (Villapol et al., 2017). Brain tissue sections were dehydrated through a series of ethanol concentrations at 50%, 70%, and twice in 100% ethanol for 5 min each. Next, they were boiled for 2 min in pretreatment 2 solution. Finally, the slides were incubated in pretreatment solution 3 (protease buffer) for 30 min before hybridization. For hybridization, sections were incubated at 40°C for 2 h with specific target probes separately: Mus musculus TNF-α (Advanced Cell Diagnostics, cat. #311081) and Mus musculus IL-1β (Advanced Cell Diagnostics, cat. #316891). Additionally, the negative control probe (Advanced Cell Diagnostics, cat. #310043) and the positive control probe (Advanced Cell Diagnostics, cat. #313911) were applied and hybridized for 2 h at 40°C. The amplification steps were performed according to the manufacturer’s instructions.

### Immunofluorescence analysis and cell death assay

Brain sections underwent immunohistochemistry, starting with three detailed 5 min washes in PBS containing 0.5% Triton X-100 (PBS-T). To avoid nonspecific binding, sections were incubated for 1 h at room temperature with 5% normal goat serum (NGS) in PBS-T. The next step was an overnight incubation at 4°C using a solution of 3% NGS in PBS-T, containing primary antibodies: anti-rabbit Iba-1 (Wako, cat. #019-19741) at 1:500 to label microglia and macrophages, and anti-mouse TERT (Novus, cat. #NB100-317) at 1:500. The following day, the sections were washed three times for 5 min each in PBS-T and then incubated with the appropriate secondary antibodies, all diluted at 1:1000, for 2 h at room temperature. The sections were rinsed three times with PBS for 5 min each, then stained with a DAPI solution in PBS to label the nuclei. Afterwards, they were rinsed thoroughly with distilled water and mounted using Fluoro-Gel and Tris Buffer mounting medium.

### Quantitative analysis of immunolabeled images

All histological images were captured by a Slideview VS200 Universal Whole Slide Imaging Scanner (Evident, USA) and a confocal imaging system (Leica Microsystems, Deerfield, IL, USA). We employed unbiased, standardized sampling methods to assess tissue regions in the cortex that exhibited positive immunoreactivity for quantitative analysis of immunolabeled sections. For proportional area measurements, microglia/macrophage Iba-1 immunoreactivity was expressed as the proportion of the target region covered by immunohistochemically stained cellular profiles. To quantify the number of Iba-1 and TERT cells in the injured cortex, we counted and imaged an average of four coronal sections from the lesion epicenter (−1.34 to −2.30 mm from Bregma) for each mouse. Double co-localization of TNF-α and IL-1β mRNA with Iba-1 staining was evaluated using z-stack acquisitions on a confocal microscope.

### Organ paraffin embedding and hematoxylin and eosin staining for *in vivo* biocompatibility

The heart, lungs, liver, spleen, and kidneys were collected and fixed in 4% paraformaldehyde for 48 h before being transferred to 70% ethanol. They were processed with a Shandon Exelsion ES Tissue Processor and then embedded in paraffin according to the manufacturer’s standard procedures. The organs were sectioned into 5 μm-thick slices. These sections underwent two 30 min dehydration steps in 95% ethanol, followed by a 1 h soak in xylene at 60-70°C, and were then embedded in paraffin for 12 h. Following dehydration, the tissues were stained with hematoxylin for 6 h at 60-70°C, rinsed with tap water, differentiated with a solution of 10% acetic acid and 85% ethanol diluted in water (twice for 2 min), and washed with tap water.

### Statistical analysis

Two-way ANOVA analyses were performed to evaluate the effects of sex (male vs. female) and treatment (Luc-LNP vs. mTERT-LNP) for the groups using Tukey’s post hoc tests. A one-way analysis of variance (ANOVA) followed by Tukey’s multiple comparison test was used for biodistribution studies. Immunofluorescence staining data, including Iba-1+ and TERT+cells, were evaluated using two-way ANOVA. Student’s t-test was used for the qPCR assay. All mice were randomly assigned to experimental groups, and the experimenters remained blinded to the treatments throughout the study. Data are expressed as the mean with the standard error of the mean (± SEM). Statistical analyses were performed using GraphPad Prism 8 (GraphPad Software, San Diego, CA, USA) for multiple groups, assuming a normal distribution of all data points. Significance levels are indicated as *p < 0.05, **p < 0.01, ***p < 0.001, and ****p < 0.0001.

## RESULTS

### Design and physicochemical characterization of mRNA-LNP formulations

We generated LNP formulations encapsulating either firefly luciferase mRNA (Luc-LNPs) or mouse TERT mRNA (mTERT-LNPs) and characterized their physicochemical properties (Figure 1). All three independently prepared batches for each formulation showed high encapsulation efficiency (>95%) by RiboGreen assay (Figure 1a,d). Dynamic light scattering revealed a narrow size distribution with mean hydrodynamic diameters of ∼90–110 nm and low polydispersity indices (PDI<0.2), indicating a uniform particle population (Figure 1b,e). Both Luc-LNPs and mTERT-LNPs exhibited slightly negative zeta potential (Figure 1c,f), consistent with reduced nonspecific protein adsorption and prolonged circulation. To evaluate *in vitro* biocompatibility, we exposed N2a murine neural cells to increasing concentrations of Luc-LNPs or mTERT-LNPs (0.5-2 µg/mL mRNA). Neither formulation affected cell viability at 24 h or 48 h compared with untreated controls (Figure 1g-j). Finally, the mTERT-LNP formulation maintained stable RNA content when stored at 4°C for up to 14 days, with no significant loss in mRNA concentration (Figure 1k), supporting its suitability for *in vivo* use.

**Figure 1.**
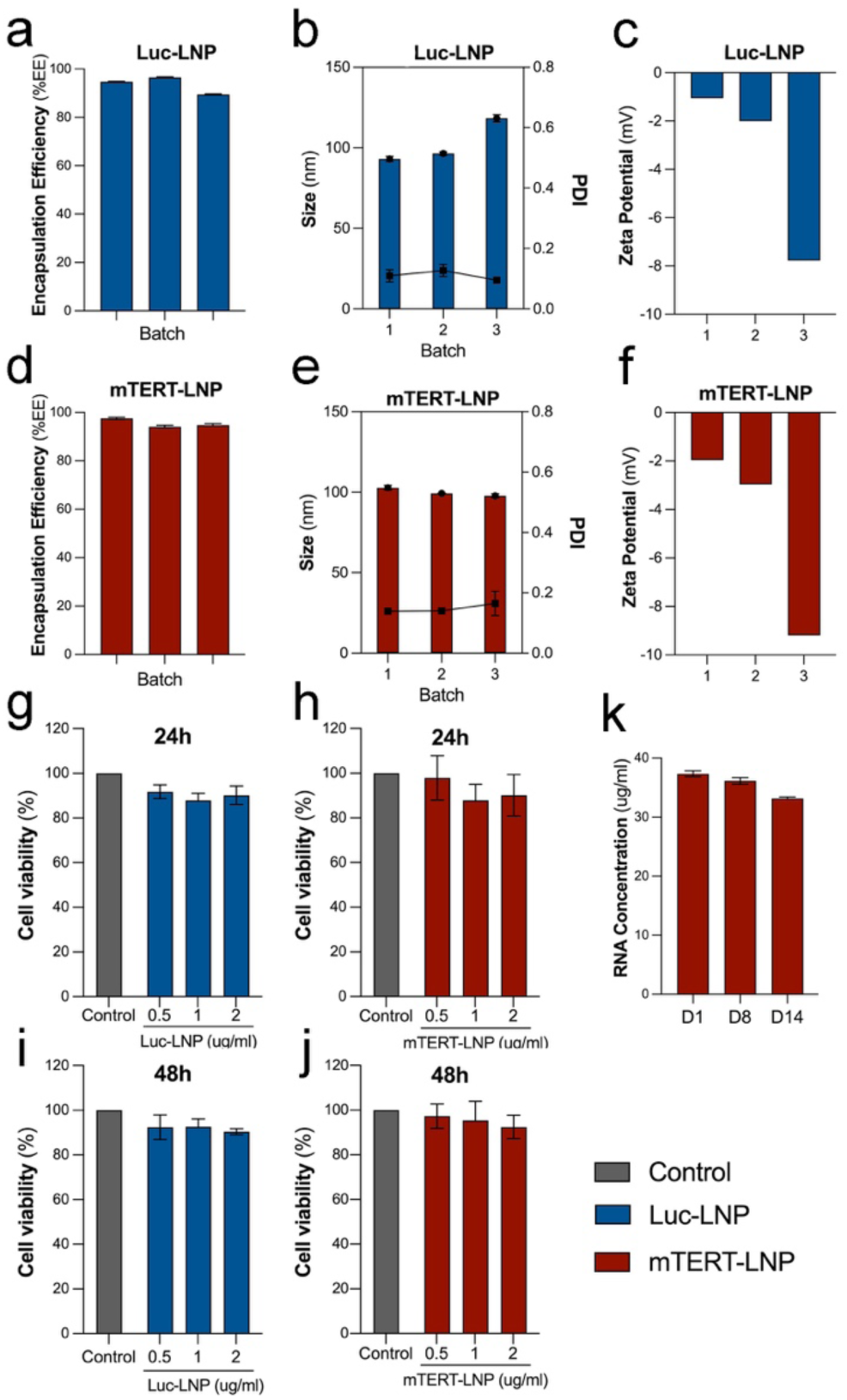
Physicochemical characterization, cytocompatibility, and storage stability of Luc- and mTERT-LNPs. (a,d) Encapsulation efficiency (%EE) of luciferase mRNA-loaded LNPs (Luc-LNP) and mouse TERT mRNA-loaded LNPs (mTERT-LNP) across three independently prepared batches. (b,e) Hydrodynamic diameter (bars, left y-axis) and polydispersity index (PDI; circles, right y-axis) of Luc-LNP (b) and mTERT-LNP (e) batches, showing uniform particle size (∼100 nm) and low PDI. (c,f) Zeta potential measurements of Luc-LNP (c) and mTERT-LNP (f) for three independent batches, indicating slightly negative surface charge. (g–j) *In vitro* cytocompatibility of Luc-LNP (g,i) and mTERT-LNP (h,j) in cultured cells at the indicated mRNA concentrations (0.5–2 µg/ml) compared with untreated control, assessed at 24 h (g,h) and 48 h (i,j). No appreciable reduction in cell viability is observed at any dose or time point. (k) mTERT-LNP RNA concentration after storage, measured on days 1, 8, and 14, demonstrating preserved RNA content over time. Data are presented as mean ± SEM of independent preparations/experiments. LNP, lipid nanoparticle; Luc, mRNA encoding luciferase; mTERT, mRNA encoding mouse telomerase reverse transcriptase; PDI, polydispersity index.

### Biodistribuon of mRNA-LNPs in brain and peripheral organs

We next evaluated the biodistribution of mRNA-LNPs using *ex vivo* IVIS imaging 24 h after intravenous administration. Mice received either Luc-LNPs or Cy5.5-labeled mTERT-LNPs (Cy5.5-mTERT-LNP). Luc-LNPs-treated animals displayed robust bioluminescent signal in the spleen and the injured brain, with minimal signal detected in the heart, lungs, kidneys, liver, or blood (Figure 2a,b). Cy5.5-LNP-mTERT showed a similar pattern of brain localization but, as expected for fluorescently labeled LNPs, also accumulated prominently in the clearance organs, including spleen, kidneys, and liver, with negligible signal in heart, lungs, or blood (Figure 2c,d). In both formulations, qualitative inspection suggested modest sex-dependent differences in organ signal intensity, with females tending to show higher signal intensity in the brain and spleen. Overall, these data confirm that IV-administered mRNA-LNPs accumulate in the injured brain and enable protein translation, while exhibiting a typical peripheral biodistribution profile for LNPs.

**Figure 2.**
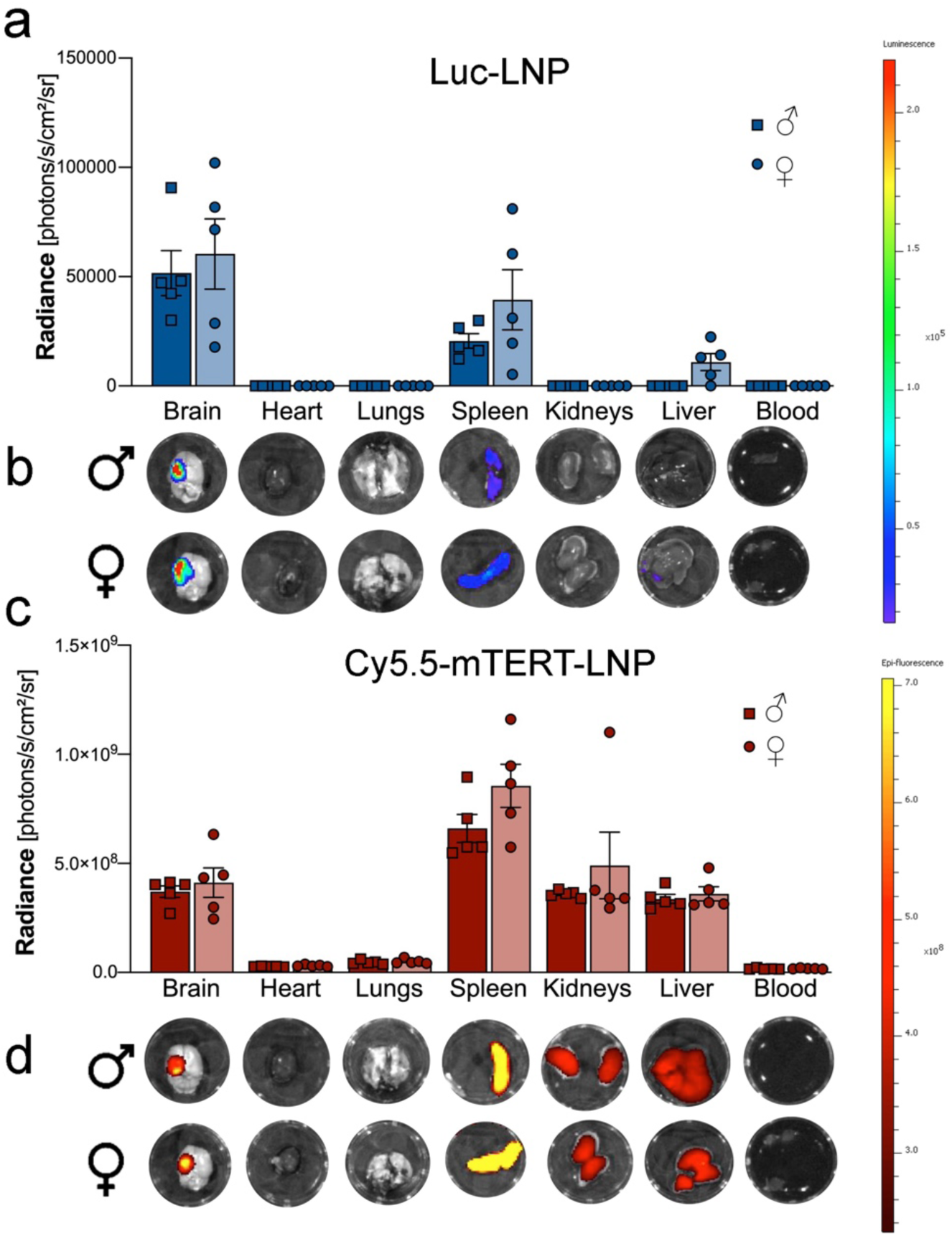
Biodistribution of systemically administered mRNA-LNPs in the injured brain and peripheral organs. (a) Quantification of bioluminescent signal (radiance, photons/s/cm²/sr) after intravenous administration of luciferase mRNA-loaded LNPs (Luc-LNP) in *ex vivo* organs collected from male (dark blue bars, square symbols) and female mice (light blue bars, square symbols) subjected to TBI. Organs analyzed include the brain, heart, lungs, spleen, kidneys, liver, and blood. (b) Representative *ex vivo* IVIS images of brain and peripheral organs from male and female mice receiving Luc-LNPs, showing luminescent signal overlaid on grayscale images. Color scale indicates relative luminescence intensity. (c) Quantification of epi-fluorescent signal (radiance, photons/s/cm²/sr) in *ex vivo* organs from male (dark red bars, square symbols) and female (light red bars, circle symbols) mice after intravenous administration of Cy5.5-labeled mTERT-LNPs (Cy5.5-LNP-mTERT). (d) Representative *ex vivo* IVIS epi-fluorescence images of brain and peripheral organs from male and female Cy5.5-LNP-mTERT-treated mice. Color scale indicates relative fluorescence intensity. Data are presented as mean ± SEM. LNP, lipid nanoparticle; Luc, mRNA encoding luciferase; mTERT, mRNA encoding mouse telomerase reverse transcriptase.

### TBI induces transient TERT/telomere dysfunction and is rescued by mTERT-LNPs

We first examined whether TBI alters endogenous TERT expression. Cortical tissue from sham-injured mice and TBI mice at 3 or 14 dpi was analyzed by qPCR. TERT mRNA levels were significantly reduced at 3 dpi and partially recovered by 14 dpi (Figure 3b). We then evaluated whether TBI affected telomere length. Telomere length, measured as the telomere-to-single-copy gene (T/S) ratio, was lower in mice at 3 dpi, but showed partial recovery by 14 dpi (Figure 3c). RNAscope confirmed a reduction of Tert mRNA signal in peri-contusional cortex at 3 dpi in both male and female mice relative to controls (Figure 3d), indicating an early window of TERT insufficiency after TBI. To test whether exogenous TERT mRNA could restore Tert expression in this window, 6-month-old male and female C57BL/6 mice received a single retro-orbital injection of mTERT-LNPs or control Luc-LNPs shortly after TBI induction (Figure 3a). At 3 dpi, RNAscope quantification showed a robust increase in cortical Tert mRNA signal in mTERT-LNPs-treated mice compared with Luc-LNPs controls in both sexes (Figure 3e,g). qPCR corroborated this finding, demonstrating significantly higher TERT mRNA levels in the mTERT-LNP group (Figure 3f). We next assessed whether TERT restoration impacted telomere length. Telomere length tended to be higher in mTERT-LNPs-treated mice compared with Luc-LNPs controls at 3 dpi, with a similar pattern in males and females (Figure 3h). Consistent with enhanced transcript levels, immunohistochemistry revealed a significant increase in the number of TERT-positive cells in peri-contusional cortex following mTERT-LNP treatment in both sexes (Figure 3i,j). Together, these data demonstrate that systemically delivered mTERT-LNPs effectively increase cortical TERT expression in the injured brain and partially restore telomere homeostasis.

**Figure 3.**
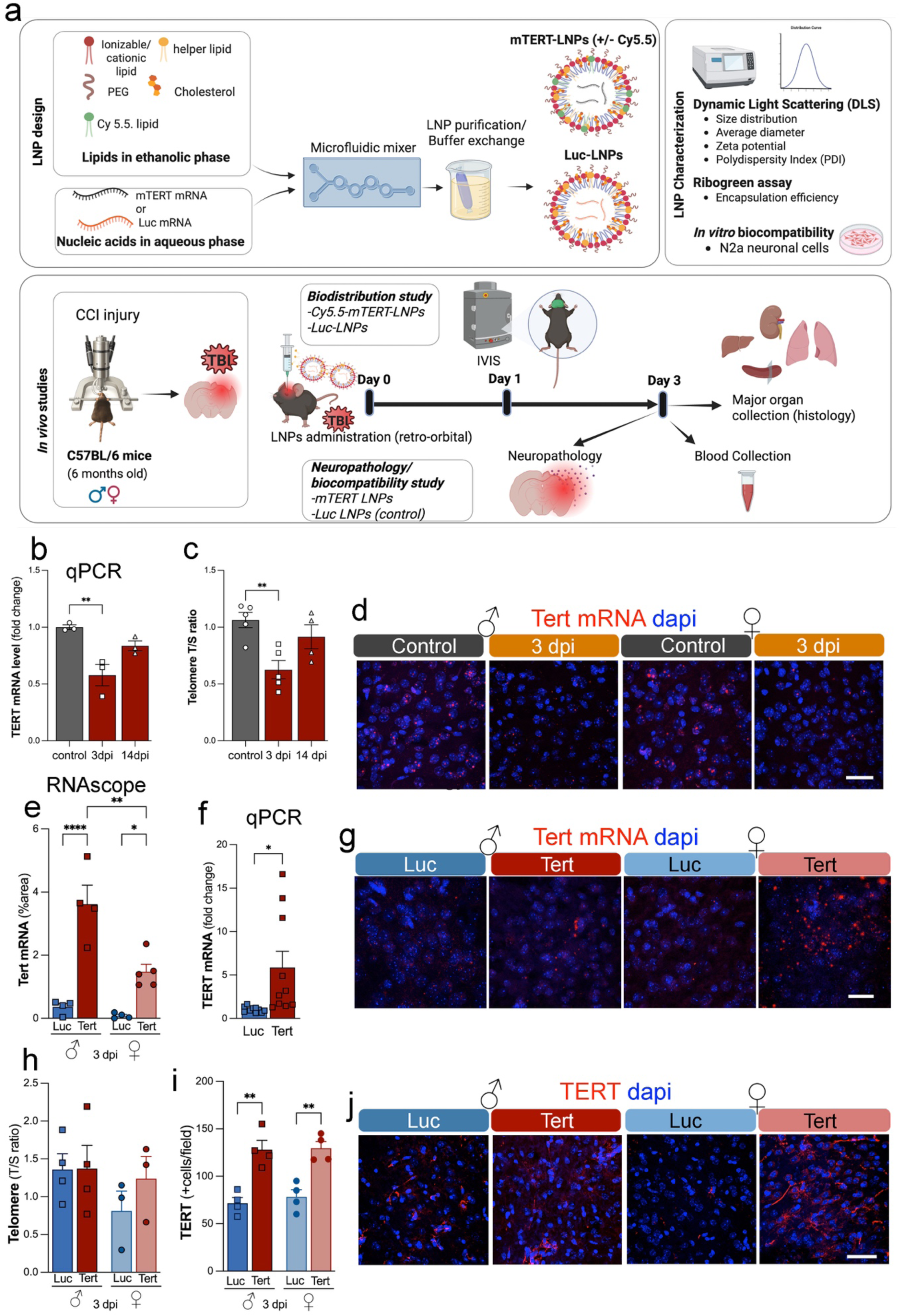
Experimental design and TERT mRNA-LNP-mediated restoration of TERT expression and telomere length after TBI. (a) Schematic of the TERT mRNA-LNP system and in vivo study design. TERT or luciferase mRNA is encapsulated in Cy5.5-labeled LNPs and administered intravenously to 6-month-old male and female C57BL/6 mice subjected to controlled cortical impact TBI (day 0). Mice undergo IVIS imaging for mRNA-LNP biodistribution and protein expression on day 1 and neuropathologic and blood analyses on day 3. (b) Endogenous brain TERT mRNA levels were assessed by qPCR at baseline (control), 3 days post-injury (dpi), and 14 dpi, demonstrating an early decrease in TERT expression with partial recovery over time. (c) The telomere length (measured by the telomere-to-single-copy gene (T/S) ratio) decreased at 3 dpi and showed partial recovery by 14 dpi. (d) Representative RNAscope images of Tert mRNA (red) with DAPI nuclear counterstain (blue) in peri-contusional cortex from male and female mice at control and 3 dpi, illustrating reduced Tert signal after TBI. Scale bar, 50 µm. (e) Quantification of RNAscope Tert mRNA signal (% area) in peri-contusional cortex from male and female mice treated with luciferase control LNPs (Luc) or TERT mRNA-LNPs (Tert) and analyzed at 3 dpi, showing robust enhancement of Tert mRNA with TERT mRNA-LNP treatment. (f) qPCR analysis of cortical Tert mRNA levels (fold change) in Luc versus Tert groups at 3 dpi, confirming increased Tert transcript levels after TERT mRNA-LNP delivery. (g) Representative RNAscope images of Tert mRNA (red) with DAPI (blue) in peri-contusional cortex of male and female mice receiving Luc or mTERT LNPs. Scale bar, 50 µm. (h) Telomere length in peri-contusional cortex measured by qPCR (T/S ratio) at 3 dpi, showing partial restoration of telomere length in Tert-treated animals relative to Luc controls, in both sexes. (i) Quantification of TERT protein+ cells per field in peri-contusional cortex by immunohistochemistry (IHC), demonstrating increased TERT protein expression in Tert versus Luc groups at 3 dpi. (j) Representative IHC images of TERT protein (red) with DAPI (blue) in cortex from male and female Luc- and Tert-treated mice. Scale bar=50 µm. Data points represent individual mice; bars show mean ± SEM. Statistical significance is indicated by *p<0.05, **p<0.01, ****p<0.0001.

### mTERT-LNPs attenuate microglial activation and pro-inflammatory cytokine expression

To determine whether TERT restoration modulates neuroinflammation, we quantified microglial activation in the peri-contusional cortex at 3 dpi. Immunohistochemistry for Iba-1 revealed that mTERT-LNP treatment significantly reduced the number of Iba-1+ microglia in both male and female mice compared with Luc-LNP controls (Figure 4a). A similar trend was observed for Iba-1–positive area, particularly in males (Figure 4b). Representative images illustrate reduced microglial density and a less activated morphology in mTERT-LNPs-treated brains (Figure 4c). We then analyzed inflammatory gene expression by qPCR. Cortical mRNA levels of the pro-inflammatory cytokines IL-1β, TNF-α, and IL-6 were significantly lower in mTERT-LNPs-treated mice than in Luc-LNPs controls, while IL-18 was unchanged (Figure 4d-f). In contrast, expression of the anti-inflammatory cytokines TGF-β and IL-10 showed a modest, non-significant increase (Figure 4g-i). These data indicate that mTERT-LNP therapy selectively suppresses key components of the acute pro-inflammatory response after TBI while sparing or slightly promoting anti-inflammatory pathways.

**Figure 4.**
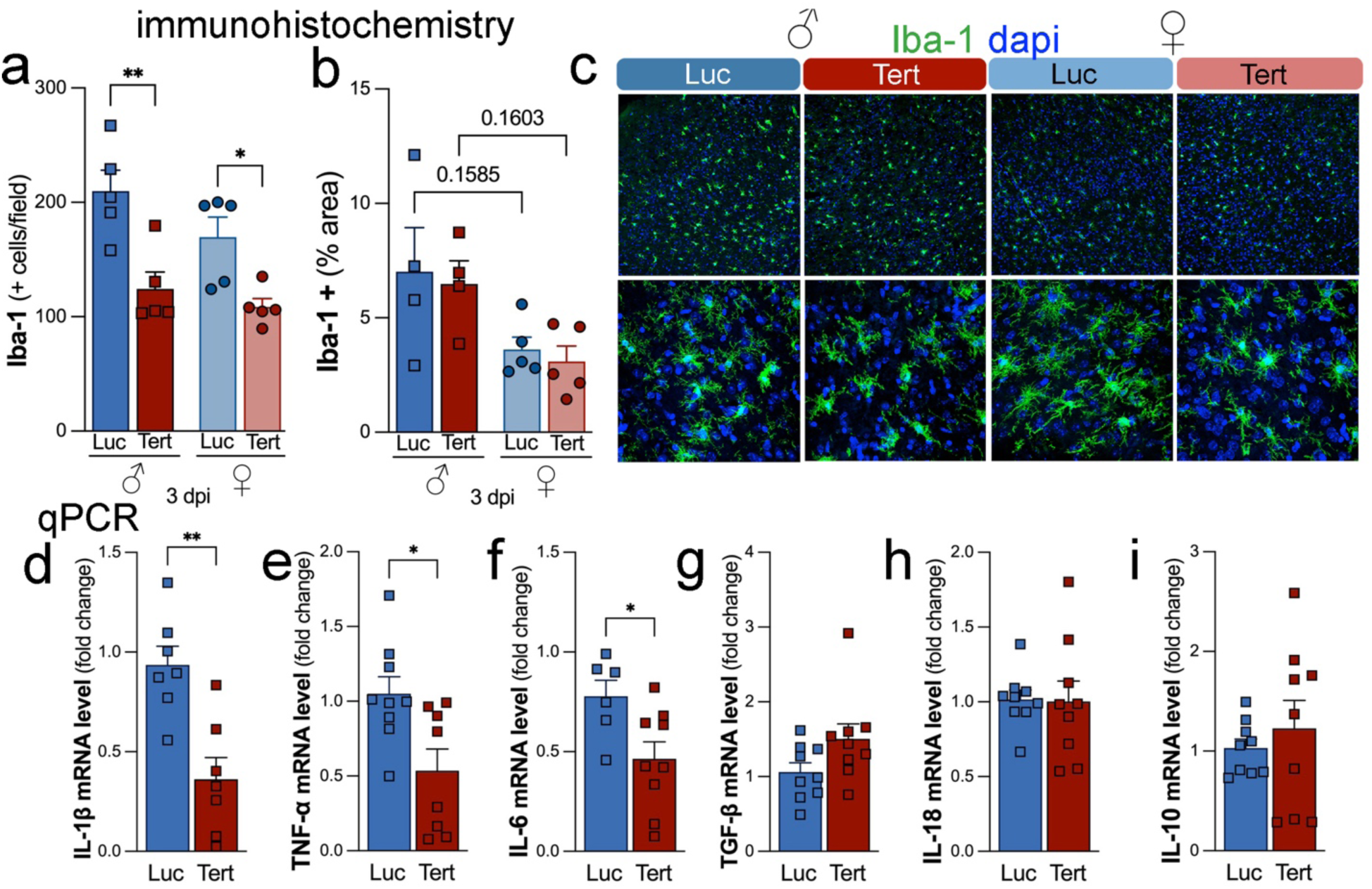
TERT mRNA-LNP treatment attenuates microglial activation and pro-inflammatory cytokine expression after TBI. (a,b) Quantification of microglial activation in peri-contusional cortex at 3 days post-injury (dpi) in male and female mice treated with luciferase control LNPs (Luc) or TERT mRNA-LNPs (Tert). Iba-1+ cells per field (a) and Iba-1+ area (%) (b) were measured by immunohistochemistry, showing a significant reduction in Iba-1+ cell number with Tert treatment in both sexes, with a trend toward reduced Iba-1+ area. (c) Representative confocal images of Iba-1 immunofluorescence (green) with DAPI nuclear counterstain (blue) in peri-contusional cortex from male and female Luc- and Tert-treated mice at 3 dpi. Top panels show low-magnification views; bottom panels show higher-magnification insets highlighting microglial morphology. (d-i) qPCR analysis of cortical cytokine mRNA expression at 3 dpi comparing Luc and Tert groups: IL-1β (d), TNF-α (e), IL-6 (f), TGF-β (g), IL-18 (h), and IL-10 (i). Tert treatment significantly reduced IL-1β, TNF-α, and IL-6 transcripts, with no significant changes in TGF-β, IL-18, or IL-10. Data points represent individual mice; bars indicate mean ± SEM. Statistical significance is denoted as *p< 0.05, **p< 0.01.

### mTERT-LNP therapy reduces peripheral inflammation and oxidative stress and is well tolerated systemically

To investigate systemic effects of TERT mRNA therapy, we measured circulating markers of inflammation and oxidative stress at 3 dpi. Western blot analysis showed that serum C-reactive protein (CRP) levels were significantly reduced in mTERT-LNP-treated male mice relative to Luc-LNP controls, with a similar but less pronounced trend in females (Figure 5a,b). In parallel, serum malondialdehyde (MDA), a marker of lipid peroxidation, was modestly decreased by mTERT-LNPs in both sexes, reaching statistical significance when comparing mTERT-LNP-treated females to Luc-LNP-treated males (Figure 5c). These findings suggest that mTERT-LNPs dampen not only central but also peripheral inflammatory and oxidative responses to TBI.

**Figure 5.**
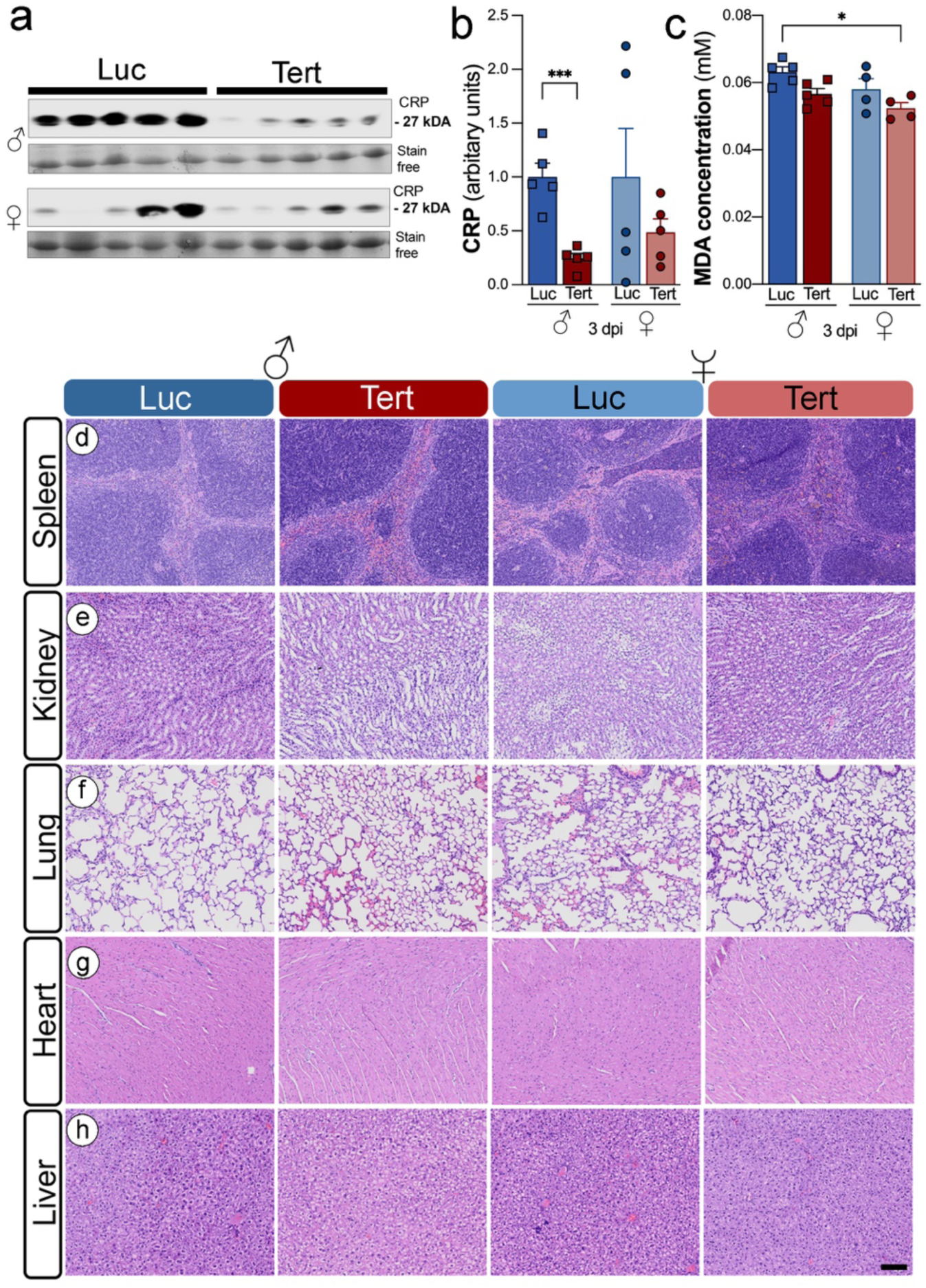
TERT mRNA-LNPs reduce systemic inflammation and oxidative stress without inducing peripheral organ toxicity. (a) Representative Western blots of serum C-reactive protein (CRP) from male and female mice at 3 days post-injury (dpi) treated with luciferase control LNPs (Luc) or TERT mRNA-LNPs (Tert). Stain-free total protein is shown as a loading control. (b) Densitometric quantification of CRP levels (arbitrary units) normalized to total protein demonstrates a significant reduction in circulating CRP in Tert-versus Luc-treated mice, particularly in males. (c) Serum malondialdehyde (MDA) concentrations, a marker of lipid peroxidation and oxidative stress, showing decreased MDA in Tert-treated animals at 3 dpi. (d–h) Representative hematoxylin and eosin (H&E)-stained sections of spleen (d), kidney (e), lung (f), heart (g), and liver (h) from male and female Luc- and Tert-treated mice at 3 dpi. No treatment-related histopathologic abnormalities are observed in any organ, indicating that TERT mRNA-LNP administration is well tolerated systemically. Scale bar in (h), 100 µm (applies to all histological panels). Data points represent individual mice; bars indicate mean ± SEM. Statistical significance: *p< 0.05, ***p< 0.001.

To assess safety, we performed histopathologic analysis of the spleen, kidney, lung, heart, and liver. Hematoxylin and eosin-stained sections from mTERT-LNP-treated mice showed no treatment-related abnormalities or tissue damage compared with Luc-LNP controls in either sex (Figure 5d–h). Consistent with this, no differences in body weight or gross behavior were observed between treatment groups (data not shown). Collectively, these results indicate that mTERT-LNP therapy exerts anti-inflammatory and antioxidant effects while maintaining a favorable systemic safety profile *in vivo*.

## DISCUSSION

This study introduces a telomerase-based nanomedicine as a therapeutic strategy for TBI. By delivering mouse TERT mRNA in a clinically inspired LNP platform, we show that a single systemic dose in the acute post-injury window efficiently reaches the injured brain, restores TERT expression, partially normalizes telomere biology, dampens microglial activation and pro-inflammatory signaling, and reduces systemic inflammatory and oxidative stress markers, all without detectable toxicity in primary organs or exacerbation of tissue damage. To our knowledge, this is the first demonstration that mRNA-LNP-mediated telomerase modulation can attenuate acute secondary injury mechanisms after TBI.

Our group has previously explored lipid-based nanocarriers as tools for diagnosis and therapy in TBI. We developed leukocyte-inspired “leukosomes,” biomimetic lipid nanoparticles that incorporate leukocyte membrane proteins into a liposomal shell and demonstrated that, after systemic administration, these particles preferentially home to inflamed vasculature in the injured brain, with minimal accumulation in uninjured cortex (Zinger et al., 2021). In a mouse CCI model, leukosomes enabled noninvasive in vivo imaging of nanoparticle trafficking to the lesion. They revealed a characteristic distribution across spleen and other clearance organs, providing a blueprint for systemic nanocarrier delivery in TBI. More recently, we extended our LNP work to mRNA cargos, showing that hTERT mRNA-LNPs enhance telomerase activity, proliferation, and engraftment of human skin cell suspensions in a wound model, establishing the feasibility and safety of transient telomerase mRNA delivery *in vivo* (Chang et al., 2024). We have also recently demonstrated that hTERT mRNA-LNPs protect from radiation-induced DNA damage in the skin (Li et al., 2025). Building on these platforms, the current study represents the first application of TERT-mRNA-LNPs directly in a neurotrauma setting. It complements ongoing efforts in our group to adapt LNP formulations for the delivery of genome-editing and other RNA therapeutics to the injured brain.

Our formulations displayed physicochemical properties consistent with state-of-the-art, clinically validated LNP systems (Wu et al., 2024). Both Luc-LNPs and mTERT-LNPs exhibited >95% encapsulation efficiency, narrow size distributions with an average diameter of ∼100 nm with low PDI, mildly negative zeta potentials, and they remained stable for at least two weeks. *In vitro*, neither formulation impaired N2a cell viability over a range of therapeutically relevant mRNA doses. These data, together with the absence of histopathologic abnormalities in spleen, kidney, lung, heart, or liver, and stable body weight *in vivo*, support the overall biocompatibility of the platform (Zhang et al., 2024).

Biodistribution studies revealed that intravenously administered mRNA-LNPs reached the injured brain while following a typical LNP clearance pattern (Bharadwaj et al., 2018). Luc-LNPs generated a strong bioluminescent signal in the brain and spleen, indicating efficient protein expression, whereas Cy5.5-labeled mTERT-LNPs showed fluorescence in the brain, spleen, kidneys, and liver, which are organs enriched in reticuloendothelial and filtration functions (Zhang et al., 2024). The discrepancy between Cy5.5 fluorescence and luciferase signal in clearance organs underscores the complexity of mRNA-LNP pharmacokinetics and pharmacodynamics: tissue exposure to intact nanoparticles does not necessarily translate into efficient endosomal escape, cytosolic mRNA release, and protein production (Huang et al., 2024). These data reinforce the need to interpret biodistribution and mRNA expression together when optimizing CNS-directed mRNA therapeutics.

A central mechanistic observation is that TBI induces a transient disruption of telomere/TERT homeostasis. We found that endogenous TERT mRNA levels in the cortex were significantly reduced at 3 days post-injury, with partial recovery by 14 days, and that TL (T/S ratio) was similarly decreased acutely and then rebounded. These findings are consistent with reports that TBI and other brain insults accelerate telomere shortening and that shorter telomeres correlate with poorer clinical outcomes and persistent symptom burden (Martha et al., 2023). Our data extend this work by identifying a discrete early window of TERT insufficiency in the injured brain, providing a rational temporal target for telomerase-based interventions.

Within this window, mTERT-LNP treatment robustly increased TERT mRNA and protein in peri-contusional cortex, accompanied by partial restoration of TL. These effects were observed in both sexes, with a tendency toward greater telomere rescue in females, echoing prior work where pharmacologic telomerase activation preferentially increased TERT expression in female RmTBI rats (Eyolfson et al., 2020). TERT appears to act primarily on secondary injury pathways, such as inflammation and oxidative stress, rather than reversing established tissue loss. This distinction aligns with its known roles in cellular stress responses and mitochondrial function rather than classical neuroprotection alone.

Neuroinflammation is a major driver of progressive pathology after TBI and a key link between acute injury and late-onset neurodegeneration (Shao et al., 2022). We found that mTERT-LNPs significantly reduced Iba-1+ microglial density in the peri-contusional cortex at 3 dpi in both males and females, indicating a broad attenuation of microglial activation. At the transcriptional level, mTERT-LNPs lowered cortical IL-1β, TNF-α, and IL-6 mRNA, while leaving IL-18 unchanged and modestly increasing IL-10 and TGF-β. This cytokine profile suggests a shift away from a strongly pro-inflammatory state toward a more balanced or pro-resolving milieu. These findings are in line with previous studies in aging, progeroid, and toxin-exposure models, where telomerase activation or hTERT mRNA delivery reduced pro-inflammatory cytokines, dampened microglial or glial activation, and improved neurogenesis and cognition (Raj et al., 2015; Li et al., 2019; Wang et al., 2024). A study reported that the TERT activator compound (TAC) increased TERT transcription, decreased the production of the proinflammatory cytokines IL-1β and IL-6, and promoted adult neurogenesis while also reducing neuroinflammation in the hippocampus of aged mice (Shim et al., 2024). Importantly, systemic administration of a TERT-activator compound restored hippocampal TERT levels, reduced the same pro-inflammatory cytokines, attenuated microglial reactivity, and ultimately improved cognitive performance (Zheng et al., 2025). Together, our data support a model in which restoring TERT expression in the injured brain disrupts non-canonical TERT functions, thereby limiting the acute neuroimmune response that seeds long-term neurodegeneration.

We also observed systemic benefits of mTERT-LNP therapy. Serum CRP, a clinical biomarker of systemic inflammation, was significantly reduced in treated males, with a similar trend in females, and circulating MDA, a marker of lipid peroxidation, was modestly decreased, reaching significance in female mice compared with male controls. These results echo epidemiological and experimental evidence that telomere attrition is associated with elevated CRP, oxidative stress, and increased cardiovascular risk (Herrmann and Herrmann, 2020). By reinforcing telomere/telomerase function, mTERT-LNPs may ameliorate a broader senescence-associated inflammatory phenotype that extends beyond the CNS. The sex-specific patterns we observed, including stronger CRP reduction in males and more pronounced MDA lowering in females, highlight the need to further dissect sex-dependent telomere and TERT biology and immune responses in TBI.

Crucially, the overall safety profile of mTERT-LNPs was favorable. Histological examination revealed no evidence of acute toxicity or structural damage in key organs. Unchanged body weight and normal behavior suggest that temporary systemic telomerase enhancement, at least over the short term and in the context of acute TBI, does not introduce overt safety liabilities. Our earlier studies show that TERT mRNA delivery enables only a transient (short-term, <72h) expression, which appreciably extends the telomere lengths, exhibits non-canonical TERT functions, and increases replicative capacity, but doesn’t induce immortalization (Ramunas et al., 2015b; Li et al., 2017; Li et al., 2019; Mojiri et al., 2021). Nevertheless, permanent overexpression of telomerase is well known for its role in oncogenesis, and long-term studies will be essential to exclude late adverse effects, especially in models with extended survival or repeated dosing.

This work has several limitations. First, we focused on a single dose and early time point (3 dpi); whether telomerase mRNA therapy confers sustained benefits at subacute or chronic stages remains unknown. Second, we did not yet evaluate functional outcomes such as cognition, motor performance, or mood, which are critical for translational relevance. Third, while we demonstrate changes in telomere length, microglial activation, and cytokine expression, the precise downstream pathways, telomere-dependent versus telomere-independent (e.g., mitochondrial, NF-κB, or DNA damage responses), through which TERT acts in the injured brain, remain to be elucidated. Finally, although our LNP formulation is inspired by clinical products, detailed dose-response, PK/PD modeling, and head-to-head comparisons with alternative delivery systems will be needed for optimization.

Future studies should, therefore, extend TERT-LNP treatment into chronic phases and assess behavioral and cognitive outcomes; dissect sex-specific mechanisms and tailor dosing to maximize benefit in males and females; interrogate telomere-independent TERT actions, particularly mitochondrial protection and modulation of DNA damage signaling; and explore combinatorial approaches, for example, coupling TERT mRNA with anti-excitotoxic or pro-synaptic agents, to achieve broader disease modification. Given that telomere attrition and chronic inflammation are shared features of multiple neurodegenerative diseases, these investigations may also inform the use of TERT mRNA-LNPs in conditions such as Alzheimer’s disease, Parkinson’s disease, or chronic traumatic encephalopathy.

## Conclusions

In summary, we provide the first *in vivo* evidence that telomerase mRNA nanotherapy mitigates acute secondary injury processes after TBI. A single systemic administration of mTERT-LNPs in the early post-injury window effectively targets the injured brain, enhances TERT expression, partially restores telomere homeostasis, and suppresses microglial activation and pro-inflammatory cytokine production. These central effects are accompanied by reduced circulating CRP and MDA, indicating attenuation of systemic inflammation and oxidative stress, while maintaining a favorable safety profile with no detectable organ toxicity. Together, these data support TERT restoration via mRNA-LNP delivery as a mechanistically grounded, scalable, and clinically compatible strategy to interrupt telomere-and TERT-driven inflammatory pathways after TBI. Given the established clinical use of LNP-based mRNA therapeutics and the broad relevance of telomere and telomerase biology to neurodegeneration, this platform holds substantial translational promise for TBI and other age- and inflammation-associated brain disorders.

## Acknowledgements

This study was supported by NIH grant R21NS106640 (S.V., B.G.) from the National Institute for Neurological Disorders and Stroke (NINDS) R56AG080920 (S.V.) from the National Institute on Aging (NIA). BG and JPC acknowledge partial support from CDMRP HT9425-24-1-0842/MB230026, RP200619 (Cancer Prevention Institute of Texas). JPC acknowledges partial support from 80ARC023CA002, NASA. The content is solely the responsibility of the authors and does not necessarily represent the official views of the NIH. A.M. is supported by a training fellowship from the Gulf Coast Consortia on the NLM Training Program in Biomedical Informatics & Data Science (T15LM007093).

